# Most calbindin-immunoreactive neurons, but few calretinin-immunoreative neurons, express the m1 acetylcholine receptor in the middle temporal visual area of the macaque monkey

**DOI:** 10.1101/271924

**Authors:** Jennifer J. Coppola, Anita A. Disney

**Affiliations:** Department of Psychology, Vanderbilt University, PMB 407817, 2301 Vanderbilt Place, Nashville, TN 37240-7817, USA

**Keywords:** primates, cerebral cortex, visual cortex, calcium-binding proteins

## Abstract

Release of the neuromodulator acetylcholine into cortical circuits supports cognition, although its precise role and mechanisms of action are not well-understood. Little is known about functional differences in cholinergic modulatory effects across cortical model systems, but anatomical evidence suggests that such differences likely exist because, for example, the expression of cholinergic receptors differs profoundly both within and between species. In the primary visual cortex (V1) of macaque monkeys, cholinergic receptors are strongly expressed by inhibitory interneurons. Here, we examine m1 muscarinic acetylcholine receptor expression by two subclasses of inhibitory interneurons—identified by their expression of the calcium-binding proteins calbindin and calretinin—in the middle temporal extrastriate area (MT) of the macaque. Using dual-immunofluorescence confocal microscopy, we find that the majority of calbindin-immunoreative neurons (55%) and only few calretinin-immunoreactive neurons (10%) express the m1 acetylcholine receptor. This differs from the pattern observed in V1 of the same species, lending further support to the notion that cholinergic modulation in cortex is tuned such that different cortical compartments will respond to acetylcholine release in different ways.

## Introduction

Acetylcholine (ACh) is a neuromodulator that contributes to the dynamic specification of cortical state (Jasper and Tessier, 1971; Metherate et al., 1992; Everitt and Robbins, 1997; Duque et al., 2000; Hasselmo and McGaughy, 2004; Steriade, 2004; Sarter et al., 2005; Herrero et al., 2008; Mena-Segovia et al., 2008). In adult primates, ACh is delivered to cortex by neurons whose cell bodies lie in the basal forebrain, and cholinergic signaling occurs partly through volume transmission (Umbriaco et al., 1994; Mrzljak et al., 1995), a signaling mode in which molecules are released from varicosities that are not apposed to a specialized receptive surface (i.e. that do not make a synapse). Under these conditions, molecules diffuse some distance through the extracellular space, eventually reaching receptors to which they can bind (Fuxe and Agnati, 1991). Volume transmission can also arise as a result of spillover from a synapse.

Because the cholinergic innervation of cortex arises from a relatively small population of subcortical neurons and ACh is then released by volume transmission, there has been an implicit assumption that cholinergic signaling is homogenous across fairly large regions in cortex. There are, however, mechanisms by which the cholinergic system can provide regionally-specific signaling. For instance, the cholinergic projection neurons of the nucleus basalis do exhibit a degree of cortical topography (Pearson et al., 1983; Price and Stern, 1983; Zaborszky et al., 2013). Furthermore, variations in the local anatomical features of cortical circuits can enhance this capacity for local control of neuromodulation.

We recently proposed the existence of distinct neuromodulatory compartments across cortex (Coppola et al., 2016). A neuromodulatory compartment is a region of tissue, defined anatomically, within which modulatory conditions are predicted to be relatively uniform and between which modulatory conditions differ profoundly. Anatomical characteristics that define compartments include patterns of axonal innervation and receptor expression, tissue tortuosity, and effectiveness of degradation pathways, among others. It is likely that these characteristics shape the way ACh interacts with cortical circuitry, and can provide a capacity for local modification of neuromodulatory inputs. Under this model, understanding the circuitry receptive to ACh, and how it differs between and within cortical areas, is critical to understanding cholinergic modulation in cortex.

The cholinergic system transduces signals through two families of receptors: nicotinic and muscarinic. Nicotinic receptors are ligand-gated cation channels, while muscarinic receptors are G protein-coupled and can act via direct channel coupling or through second messenger systems. Here, we investigate the expression of one of the five types of muscarinic receptor—the m1 ACh receptor (m1AChR), which is the muscarinic receptor subtype most strongly expressed in primate cortex (Mrzljak et al., 1993). Specifically, we investigate m1AChR expression by putatively inhibitory interneurons identified by their expression of the calcium-binding proteins calbindin-D28k (CB) and calretinin (CR). These two calcium-binding proteins, along with a third, parvalbumin (PV), are commonly used as population markers for subclasses of inhibitory interneurons in primates (Van Brederode et al., 1990; DeFelipe, 1997; Meskenaite, 1997; Zaitsev et al., 2005; Disney and Aoki, 2008; Povysheva et al., 2008). Here, we quantify m1AChR expression by CB-and CR-immunoreactive (ir) neurons in the middle temporal extrastriate visual area MT. Area MT in macaques (also known as visual area V5) contains neurons whose receptive fields are selective for the linear direction of visual motion (Maunsell and Van Essen, 1983; Britten et al., 1993) and is considered part of the dorsal (or ‘where’) pathway in vision (Mishkin and Ungerleider, 1982).

We have previously described m1AChR expression by inhibitory interneurons in macaque visual area V1. Specifically, in V1, m1AChRs are expressed by approximately 80% of PV-ir neurons, 60% of CB-ir neurons, and 40% of CR-ir neurons (Disney and Aoki, 2008). As in V1, m1AChRs are expressed by most PV-ir neurons (75%) in macaque MT (Disney et al., 2014). The goal of the present study was to complete our quantification and comparative anatomical assessment of m1AChR expression by inhibitory neurons in macaque MT by investigating expression by CB- and CR-ir neurons. We find that while CB-ir neurons in MT show similar levels of m1AChR expression to that observed in V1, expression by the CR-ir population differs, with very few CR- ir neurons in MT expressing m1AChRs.

## Materials and Methods

### Animals

Tissue was obtained from three adult male macaque monkeys (two *Macaca mulatta* and one *Macaca nemestrina*) that had been previously used in unrelated electrophysiology studies. For details of the standard protocols for the donor labs, see Oristaglio et al. (2006) and Nauhaus et al. (2012). All procedures were approved by the Institutional Animal Care and Use Committee and performed in accordance with National Institutes of Health and institutional guidelines for the care and use of animals.

### Histological preparation

Animals were euthanized by intravenous injection of sodium pentobarbital (60 mg/kg). Following abolition of pedal and corneal reflexes, animals were transcardially perfused with 0.01 M phosphate-buffered saline (PBS, pH 7.4) followed by 4 L of chilled 4% paraformaldehyde (PFA) in 0.1 M phosphate buffer (pH 7.4). The fixative was run for at least 40 minutes. The brain was then removed and blocked as necessary to provide donor labs with tissue for their needs. The remaining tissue was post-fixed over night at 4 °C in 4% PFA. The following day, the brain was transferred to 30% sucrose in PBS and stored at 4° C until it sank.

Hemispheres to be sectioned from two brains were blocked in approximately the coronal plane at the level of the lunate sulcus (with the entire lunate sulcus contained in the block) and at the anterior tip of the intraparietal sulcus. For the third brain, hemispheres were blocked in approximately the coronal plane, but at the anterior tip of the intraparietal sulcus, such that the entirety of primary visual cortex was contained within the block. The tissue from these blocks was sectioned at a thickness of 50 µm on a freezing microtome. Two 1-in-6 series of sections were set aside to provide reference sections for determining laminar and areal boundaries (Nissl and Gallyas silver stains). Remaining sections were stored at 4° C in PBS with 0.05% sodium azide.

### Antibody characterization

See Table 1 for a summary of antibodies used in this study. The polyclonal primary antibody used to detect m1AChRs was raised in rabbit against amino acids 227-353 of the human m1AChR (anti-m1; Alomone Labs, Jerusalem, Israel Cat# AMR-001 RRID:AB_2039993, lot# AN-05). An antibody directed against the same epitope (but from another vendor) passed both Western blot and preadsorption controls in macaque cortical tissue (Disney et al., 2006) and the specific lot number used in this study also passed preadsorption controls in macaque (Disney et al., 2014; Disney and Reynolds, 2014). The primary antibodies used to detect the calcium-binding proteins were obtained from Swant (Bellinzona, Switzerland). We used a monoclonal mouse anti-calbindin-D28k antibody raised against CB purified from chicken gut (anti-CB; Cat# 300 RRID:AB_10000347, lot# 18F), and a polyclonal goat anti-calretinin antibody raised against human recombinant CR (anti-CR; Cat# CG1 RRID:AB_10000342, lot# 1§.1). These antibodies have been characterized previously (Celio et al., 1990; Schwaller et al., 1999) and the specific lots used in this study passed preadsorption controls (Disney and Aoki, 2008).

**Table 1:**
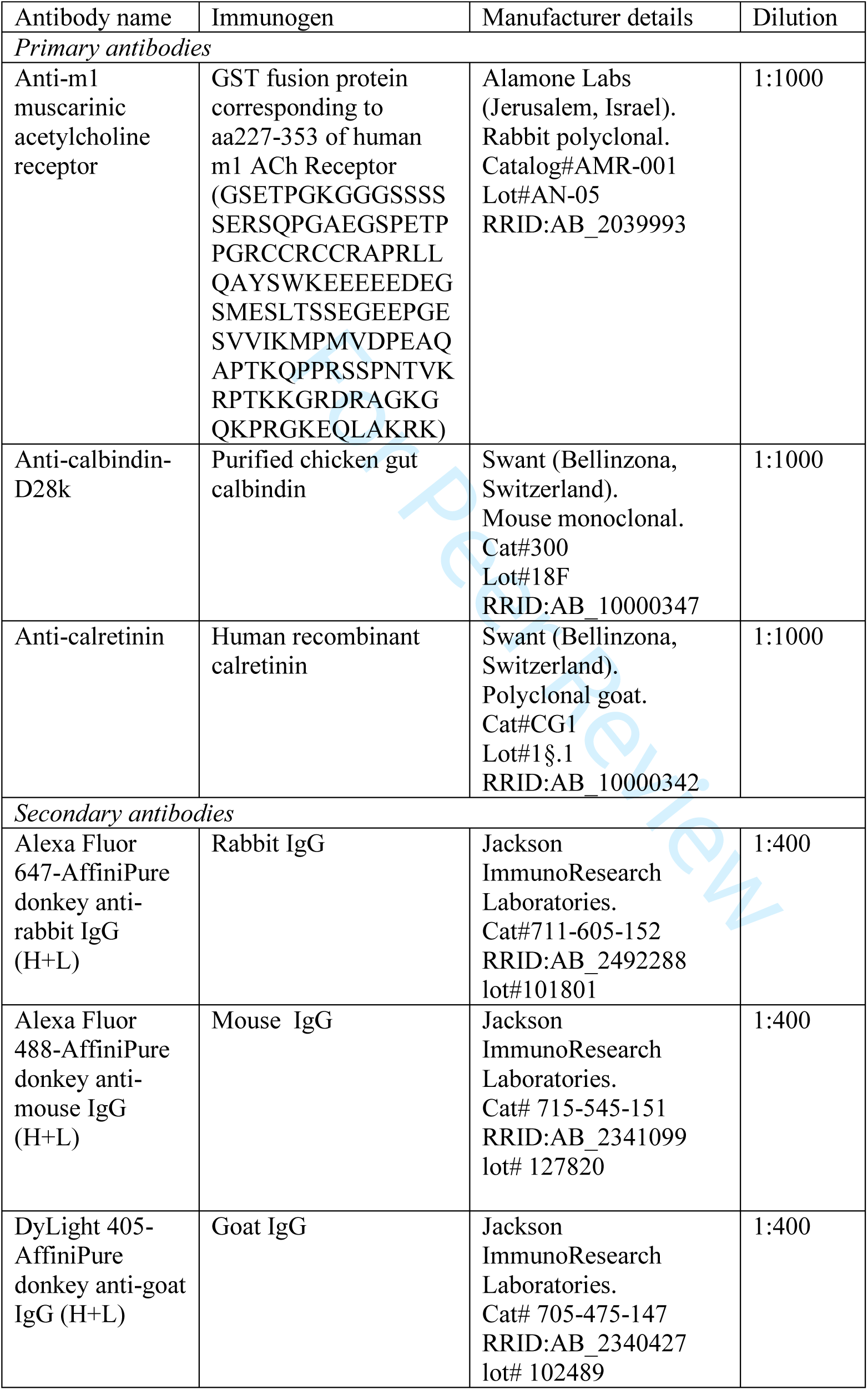
Antibody details.

F’ab fragment secondary antibodies were purchased from Jackson ImmunoResearch. The anti-m1 was detected using a donkey anti-rabbit IgG conjugated to Alexa Fluor 647 (Cat# 711-605-152 RRID:AB_2492288, lot# 101801), the anti-CB using a donkey anti-mouse IgG conjugated to Alexa Fluor 488 (Cat# 715-545-151 RRID:AB_2341099, lot# 127820), and the anti-CR using a donkey anti-goat IgG conjugated to Dylight 405 (Cat# 705-475-147 RRID:AB_2340427, lot# 102489). Reconstituted secondary antibodies were diluted 1:1 in glycerol for storage at −20°C; reported concentrations below account for this dilution. Specificity of secondary antibodies was tested by processing tissue according to the protocol below, but without any primary antibodies (no primary control). This procedure produced no fluorescent signal for any of the secondary antibodies.

### Immunohistochemistry

Dual-immunofluorescence labeling was used to visualize m1AChR immunoreactivity with immunoreactivity for either CB or CR. Two coronal sections containing area MT from each animal were chosen pseudo-randomly for each dual-labeling experiment. This resulted in 12 sections total (six sections for the CB experiment, six sections for the CR experiment). First, the tissue incubated in a 0.01M PBS blocking solution containing 1% IgG-free bovine serum albumin (BSA; Jackson ImmunoResearch), 0.05% sodium azide, and 0.5% Triton X-100 (Triton) for one hour. Next, tissue sections co-incubated at room temperature on a shaker for 72 hours in both primary antibodies at once (i.e. anti-CB and anti-m1, or anti-CR and anti-m1), each diluted at 1:1,000 in the blocking solution described above. Following the primary antibody incubation, sections were rinsed in PBS three times for 10 minutes each and then incubated for four to six hours in fluorophore-conjugated F’ab fragment secondary antibodies in the dark at room temperature on a shaker. Secondary antibodies were diluted 1:400 in PBS containing 1% BSA. Following this final incubation, sections were rinsed in PBS three times for five minutes each, mounted on subbed slides, and dried overnight. The following day, sections were dehydrated through a series of graded alcohols (50-100%), followed by 2 × 100% xylene, and coverslipped (DPX mounting medium; Sigma-Aldrich, St. Louis, MO).

### Confocal microscopy

A Zeiss LSM 710 META inverted confocal microscope driven by the Zeiss ZEN imaging software (Thornwood, NY) was used to image immunolabeled sections. The 405, 488, and 633 nm laser lines were used for fluorophore excitation with a laser power of 1.5, 2, and 10%, respectively. Laser lines were checked for “bleed-through” by independently turning off a given line and determining that no image was captured in the corresponding data channel. The pinhole was set to 61.2 µm and data for both channels collected concurrently. To determine the appropriate imaging depth within the tissue, “z-stacks” spanning the entire tissue thickness were collected just below the pial surface (i.e. in the upper portion of cortical layer II) and above the white matter (i.e. in the lower portion of cortical layer VI) in the same cortical column at 63x magnification (oil immersion). Using these z-stacks, a single imaging plane was determined at which to capture a “tilescan.” Each tilescan represents an approximately 270 µm-wide column of tissue that spans the cortex from pia to white matter and was collected at 63x (oil immersion) with 4× averaging. One tilescan was collected per tissue section and stored for off-line analysis.

### Defining architectonic boundaries

Area MT was identified using reference sections (adjacent to the immunolabeled sections) that had been stained to visualize myelin according to the Gallyas (silver) method (Gallyas, 1970). MT is located on the posterior bank of the superior temporal sulcus and is characterized by heavy myelination that has a matted appearance (Figure 1). Atlases and published data were used alongside the reference sections to identify borders at 5× magnification (Van Essen et al., 1981; Ungerleider and Desimone, 1986; Paxinos et al., 2000; Saleem and Logothetis, 2012). For each immunolabeled section, an adjacent Nissl (cresyl violet) reference section was used to identify laminar boundaries. Nissl reference section images were collected using a Carl Zeiss Axio Imager M2 light microscope with a 20× objective. Co-registration of the light and fluorescence micrographs was achieved using fiduciary marks such as pial surface shape, structural morphology, blood vessels, and cutting artifacts. To correct for differences in tissue shrinkage arising from differences in the Nissl and immunofluorescent tissue processing protocols, the distance from the pial surface to the layer I/II border (defined by a sharp increase in cell density) was measured and compared between each Nissl and tilescan image using Adobe Photoshop CS6 (San Jose, CA). Distances from the pial surface to each layer boundary (layers II/III, IV, V, VI) were measured on the Nissl section and converted to the distance for the corresponding tilescan. Laminar boundaries were drawn onto the tilescans before quantification.

**Figure 1:**
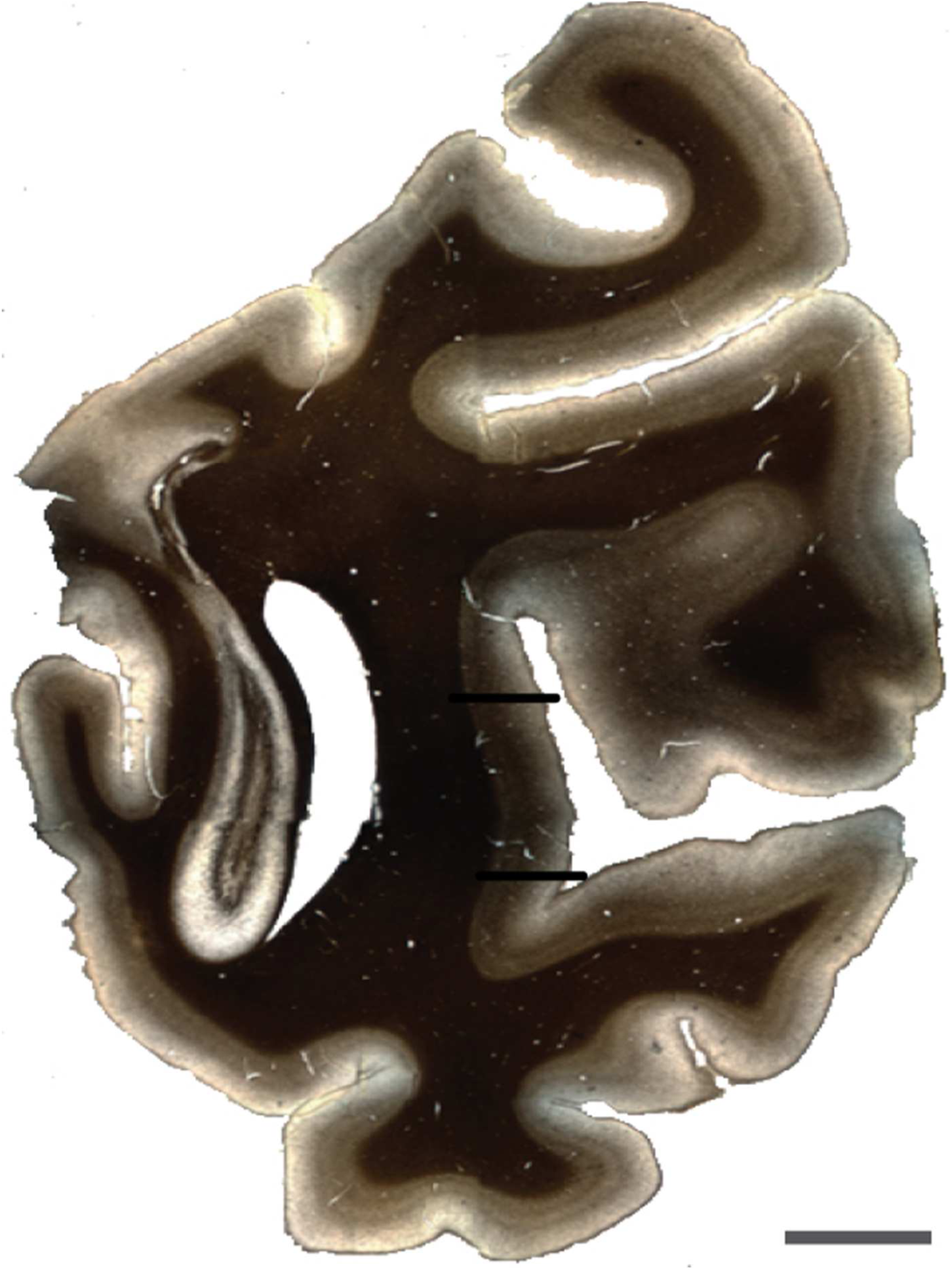
Coronal section stained to visualize myelin by the Gallyas (silver) method. Black bars show the approximate locations of the architectonic boundaries of area MT (in this section). Scale bar = 10 mm.

### Cell counts

Our goal was to determine the proportion of CB-and CR-ir neurons that express the m1AChR. Because we are interested in investigating proportional data, and not in estimating absolute cell numbers, we used a non-stereological counting method. Immunolabeled cell bodies were counted manually by a single, trained human observer using Adobe Photoshop. Each data channel was isolated and labeled cell bodies were counted in gray-scale. Only wholly visible cell bodies—with roughly 75% of the membrane in focus (determined qualitatively)—were counted. Cell bodies that touched the left boundary of the image or a laminar boundary line were not counted. In each isolated channel, immunolabeled cells that met these counting criteria were marked with shapes that reflect the size of the cell body in a separate Photoshop image layer. Dually-labeled cells were quantified by the overlap of their shape markers (Figure 2).

**Figure 2:**
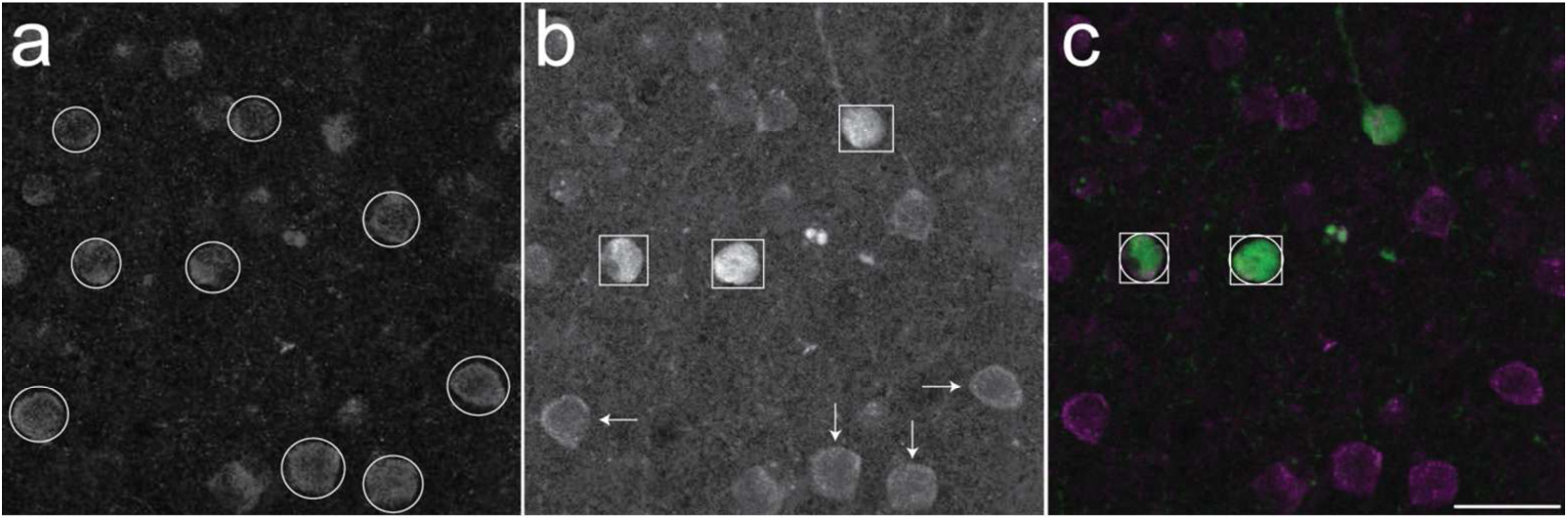
Quantification methods. (a) shows m1 acetylcholine receptor immunoreactivity. Cells marked with a circle met counting criteria and were counted. (b) shows immunoreactivity for calbindin-D28k. Within this population are two sub-groups: cells with darkly stained somata (marked with squares), and cells with faintly stained somata (marked with arrows). Only darkly stained cells were quantified. The merged imaged in (c) shows cells that were counted in both single-label images, and thus are counted as dual-labeled cells. Scale bar = 25 µm (all panels).

### Photomicrograph production

Light micrographs (of Nissl/Gallyas reference sections) were captured using a Carl Zeiss AxioCam MRc camera driven by the Stereo Investigator software package (MBF Bioscience, Williston, VT). Settings for the contrast and brightness of light micrographs were set to match the image as it appeared through the ocular lenses. Settings for contrast and brightness of the confocal images (captured using the LSM 710 confocal microscope, described above) were independently chosen for each field of view before data were collected. The conversion of data channels from red/green to magenta/green and image cropping are the only alterations made to the original confocal images for publication, with the exception of panel c in Figure 4, Figure 5, and Figure 7. For these images, gamma correction and exposure adjustments were made in Adobe Photoshop. These alterations were for publication only; all data were collected from raw, uncorrected images. No alterations were made to the light micrograph for publication.

**Figure 3:**
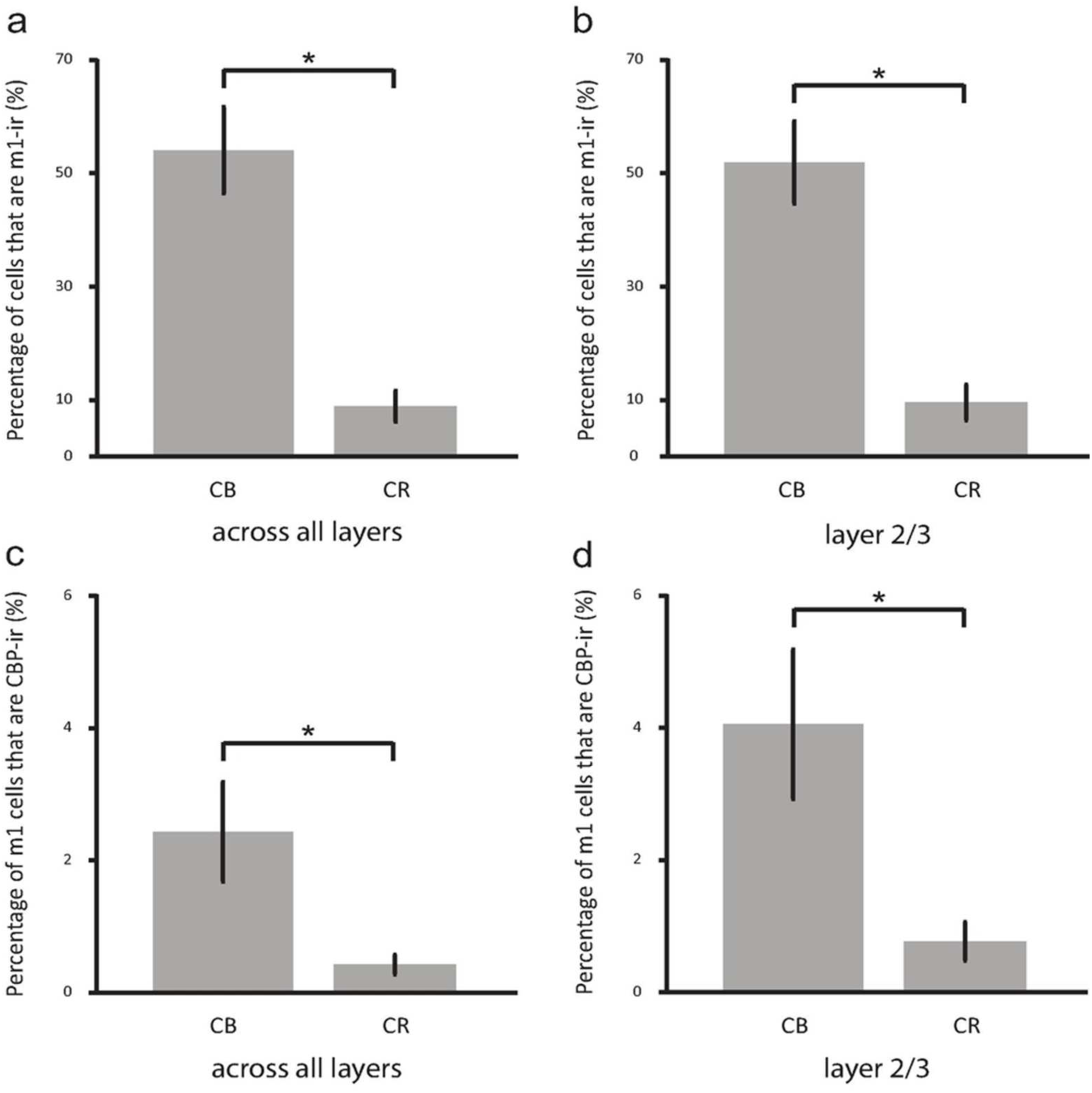
(a) shows the percentage of the calbindin-D28k (CB) and calretinin (CR) immunoreactive populations that are also immunoreactive (ir) for the m1 acetylcholine receptor (m1AChR), collapsed across all cortical layers. 55% of CB-ir neurons express m1AChRs, while 10% of CR-ir neurons express m1AChRs.(b)shows the same data (52% of CB-ir neurons and 11% of CR-ir neurons), but for layers II and III only.(c)shows the percentage of cells immunoreactive for m1AChRs that express CB or CR collapsed across all cortical layers. 2% of m1-ir neurons express CB, while 0.5% of m1-ir cells express CR. (d) shows the samedata (4% of m1AChR-ir neurons express CB, while 1% express CR), but for layers II and III only. Bars indicate standard error and asterisks indicate statistical significance (p<0.01).

**Figure 4:**
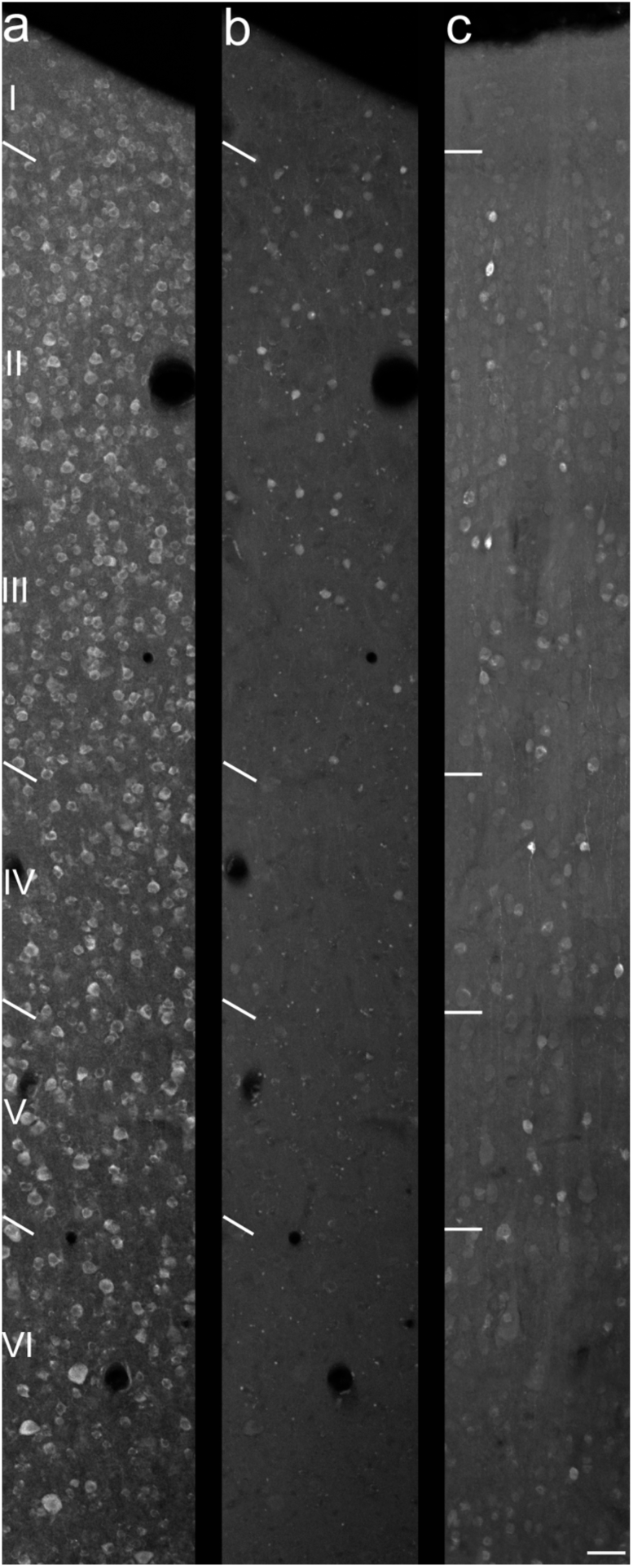
Laminar distributions of cells immunoreactive for the m1 acetylcholine receptor (a), calbindin-D28k (b), and calretinin (c). Layer boundaries are marked to the left of each panel. m1-immunoreactive neurons are more uniformly distributed throughout the tissue than are CB- and CR-immunoreactive neurons, which are mostly expressed in layers II and III. Scale bar = 50 µm (all panels).

**Figure 5:**
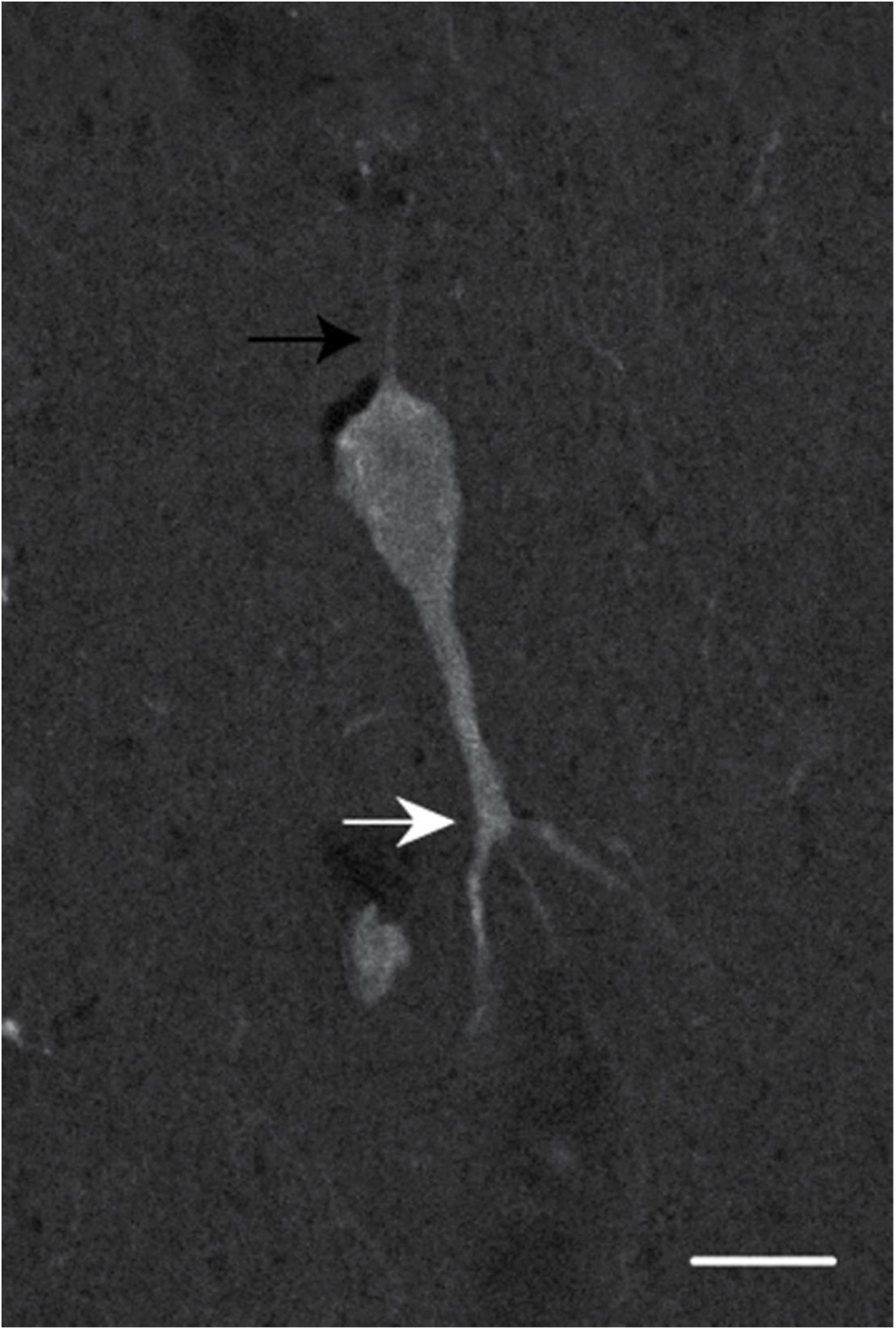
Large neuron immunoreactive for calbindin-D28k in layer V. Black arrow marks a thin axon extending out of the cell body toward the pial surface. White arrow marks the first branch point of a thick primary dendrite descending out of the cell body toward the white matter. Scale bar = 10 µm.

### Statistical analysis

Custom MATLAB (Mathworks) scripts were used for statistical analysis. Comparisons between animals revealed no significant differences. As such, all analyses combine data across all animals. Proportional testing was carried out using a two-tailed difference test on z-scores, with the following formula:

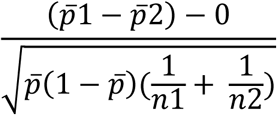

where p1 represents the observation from the first population and p2 represents the observation from the second population, and n1 denotes the total number of observations in the first population and n2 denotes the total number of observations in the second population.

## Results

We quantified dual immunofluorescence to determine the extent to which the m1AChR is expressed by putatively inhibitory interneurons that are immunoreactive for either CB or CR in area MT of the macaque monkey. Altogether, we counted 2831 immunoreactive neurons in a total area of approximately 5.4 mm^2^ across three animals. We found that 55% of CB-ir neurons express the m1AChR, while 10% of CR-ir neurons express the same receptor. Raw cell counts are listed in Table 2.

**Table 2:** Raw cell counts. The upper half of this table shows cells counted in tissue stained to visualize calbindin-D28k and m1 acetylcholine receptors, the lower half shows cells counted in tissue stained to visualize calretinin and m1 acetylcholine receptors.

| **Calbindin** |  | Layer I | Layer II/III | Layer IV | Layer V | Layer VI | Total |
| --- | --- | --- | --- | --- | --- | --- | --- |
|  | m1-ir | 64 | 689 | 283 | 228 | 187 | 1451 |
|  | CB-ir | 0 | 53 | 4 | 3 | 2 | 62 |
|  | dual-labeled | 0 | 28 | 2 | 2 | 2 | 34 |
| **Calretinin** |  |  |  |  |  |  |  |
|  | m1-ir | 30 | 717 | 231 | 159 | 121 | 1258 |
|  | CR-ir | 1 | 56 | 2 | 0 | 1 | 60 |
|  | dual-labeled | 0 | 6 | 0 | 0 | 0 | 6 |

### m1AChR expression by CB- and CR-ir neurons

Dual immunofluorescence for m1AChRs and CB/CR was found in layers II-VI. In total, a significantly higher percentage (55%) of CB-ir neurons express m1AChRs than do CR-ir neurons (10%; p<0.01, Figure 3a). As CB- and CR-ir populations are most dense within layers II and III, and are sparser in other layers (Figure 4), a statistical analysis of data from just these layers is similar to that for data collapsed across all layers. The difference in dual immunoreactivity between CB- and CR-ir populations in layers II and III considered alone is also statistically significant (p<0.01, Figure 3b).

### Calcium-binding protein expression by m1AChR-ir neurons

Across all layers, 2% of m1AChR-ir neurons express CB, and 0.5% express CR (Figure 3c). This is similar to the layer II/III data, which show 4% of m1AChR-ir express CB, while 0.8% of m1AChR-ir neurons express CR (Figure 3d). The difference in CB and CR expression by m1AChR-ir neurons is statistically significant across all layers and in layers II and III considered in isolation (p<0.01).

### Calcium-binding protein immunoreactivity

CB- and CR-ir neurons are found in all cortical layers and are most dense in layers II and III (Figure 4). This is consistent with previous reports in macaque V1 (Van Brederode et al., 1990; Meskenaite, 1997; Disney and Aoki, 2008) and MT (Dhar et al., 2001). CB immunoreactivity is exhibited by two populations of neurons interspersed within the same tissue sections and layers. One has darkly stained cell bodies while the other has lightly stained cell bodies (Figure 2). Only darkly stained neurons were quantified in this study, as these correspond to the inhibitory population (Van Brederode et al., 1990; DeFelipe, 1997; Hof et al., 1999; Disney and Aoki, 2008). Overall, few CB-ir processes were observed and morphological identification was usually not possible. However, we did observe a cell in layer V (Figure 5) that may correspond to an infragranular CB population described in V1 by Van Brederode et al. (1990). These cells are among the largest of the CB-ir neurons and resemble the type 5B:6-1 neurons described by Lund et al. (1988).

Similar to the CB-ir neurons, the CR-ir population also contains both darkly stained and lightly stained neurons. However, in the case of CR-ir neurons, a third population is also present. This type of staining is concentrated in the nucleus of the cell and in some instances, a dimmer nucleolus can be identified (Figure 6). Only darkly stained cells were quantified in this study, as these correspond to the inhibitory population (DeFelipe, 1997; Hof et al., 1999). The nuclear staining is likely a state-dependent immunoreactivity, and does not define a cell class (see Discussion). CR-ir neurons in layers II and III were often observed to be bipolar with vertically oriented processes extending from the cell body, including axonal varicosities with the classic “beads on a string” appearance (Figure 6). Of these neurons with vertically-oriented processes (Figure 7), some may correspond to the 2/3A variety 2 columnar cells described in Lund and Wu (1997). The dendrites of these neurons form a narrow columnar field, with a thin axon ascending toward the pial surface.

**Figure 6:**
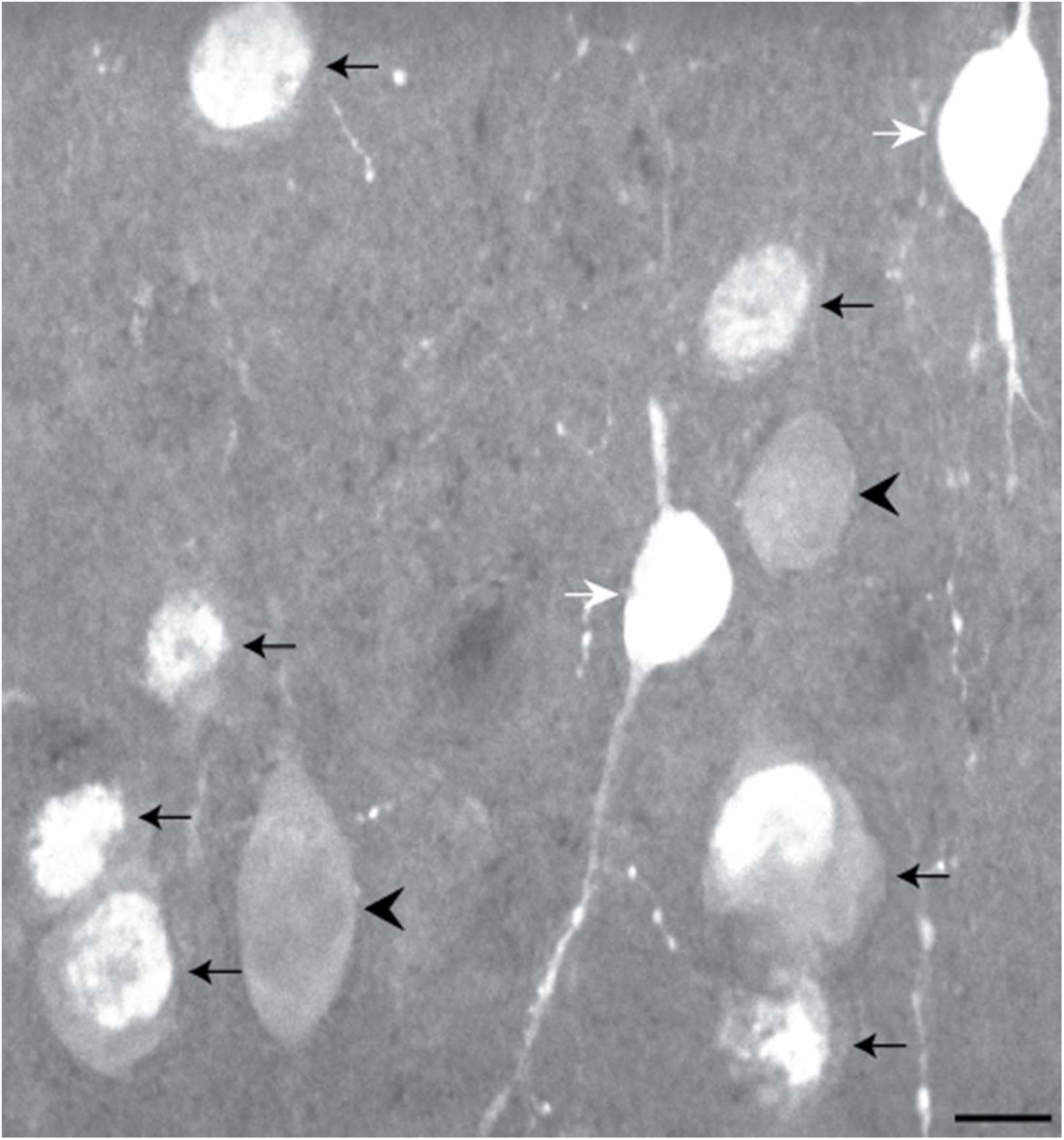
Neurons immunoreactive for calretinin in layer II. White arrows mark two cells immunoreactive for calretinin that meet the counting criteria and were quantified. From these cell bodies extend vertically oriented processes. Black arrows mark cells that show immunoreactivity within what appears to be the nucleus of the cell. In some cases, a dimmer nucleolus is also visible. Cells showing only nuclear immunoreactivity such as this were not counted. Arrowheads mark cells that are considered lightly stained and were also not counted. Scale bar = 10 µm.

**Figure 7:**
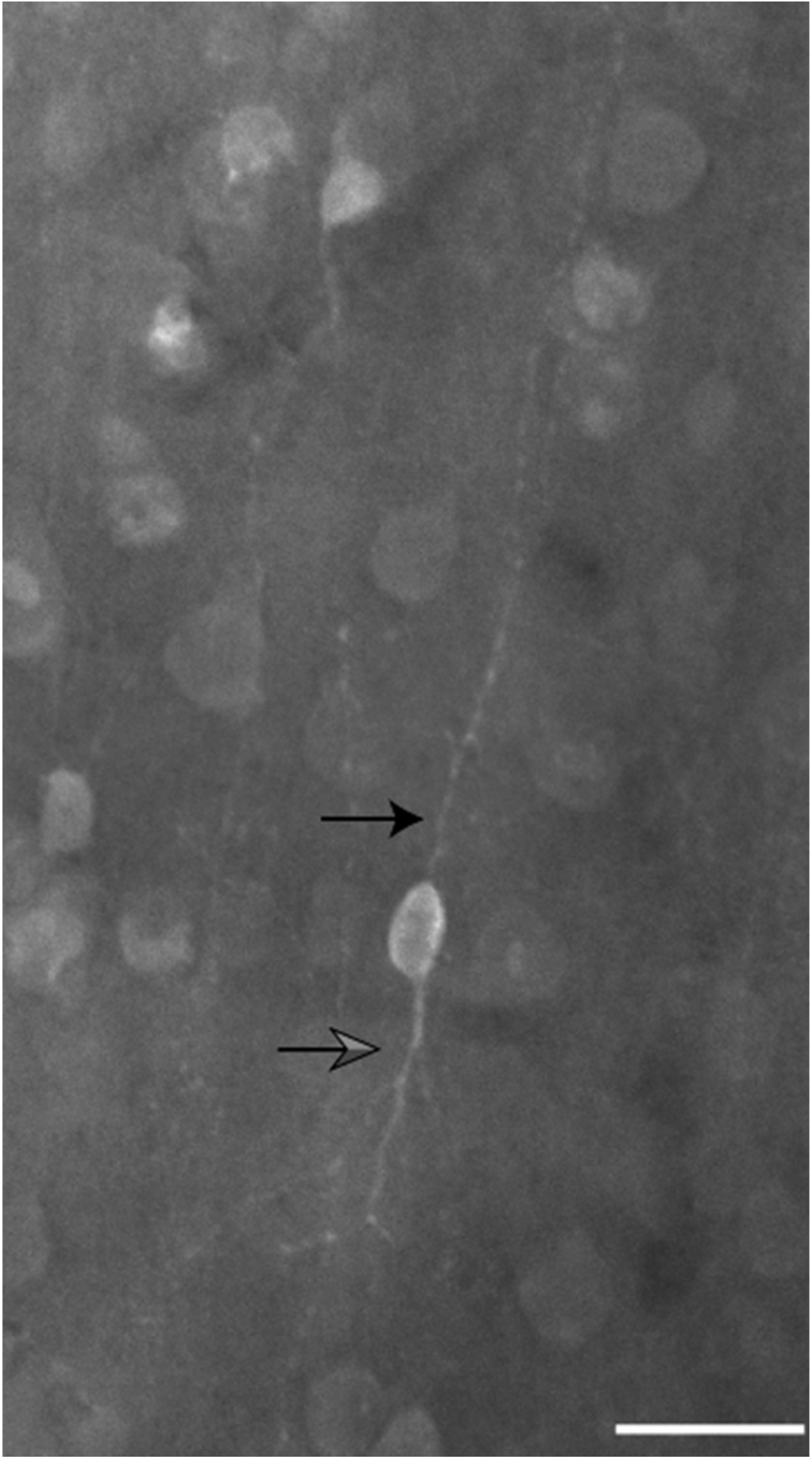
Bipolar neuron immunoreactive for calretinin in layer III. Black arrow marks a thin, varicose axon extending toward the pial surface. Gray arrow marks a primary dendrite descending from the cell body toward the white matter. Scale bar = 25 µm.

### m1AChR immunoreactivity

m1AChR-ir neurons are found in all layers. They are characterized by strong somatic labeling, which appears more intense along the perimeter of the cell body, with a more dimly stained cytoplasm (Figure 8). While few processes are visible, many that are visible are connected to a cell body. This is consistent with previous reports (Disney et al., 2006; Disney and Aoki, 2008; Disney and Reynolds, 2014).

**Figure 8:**
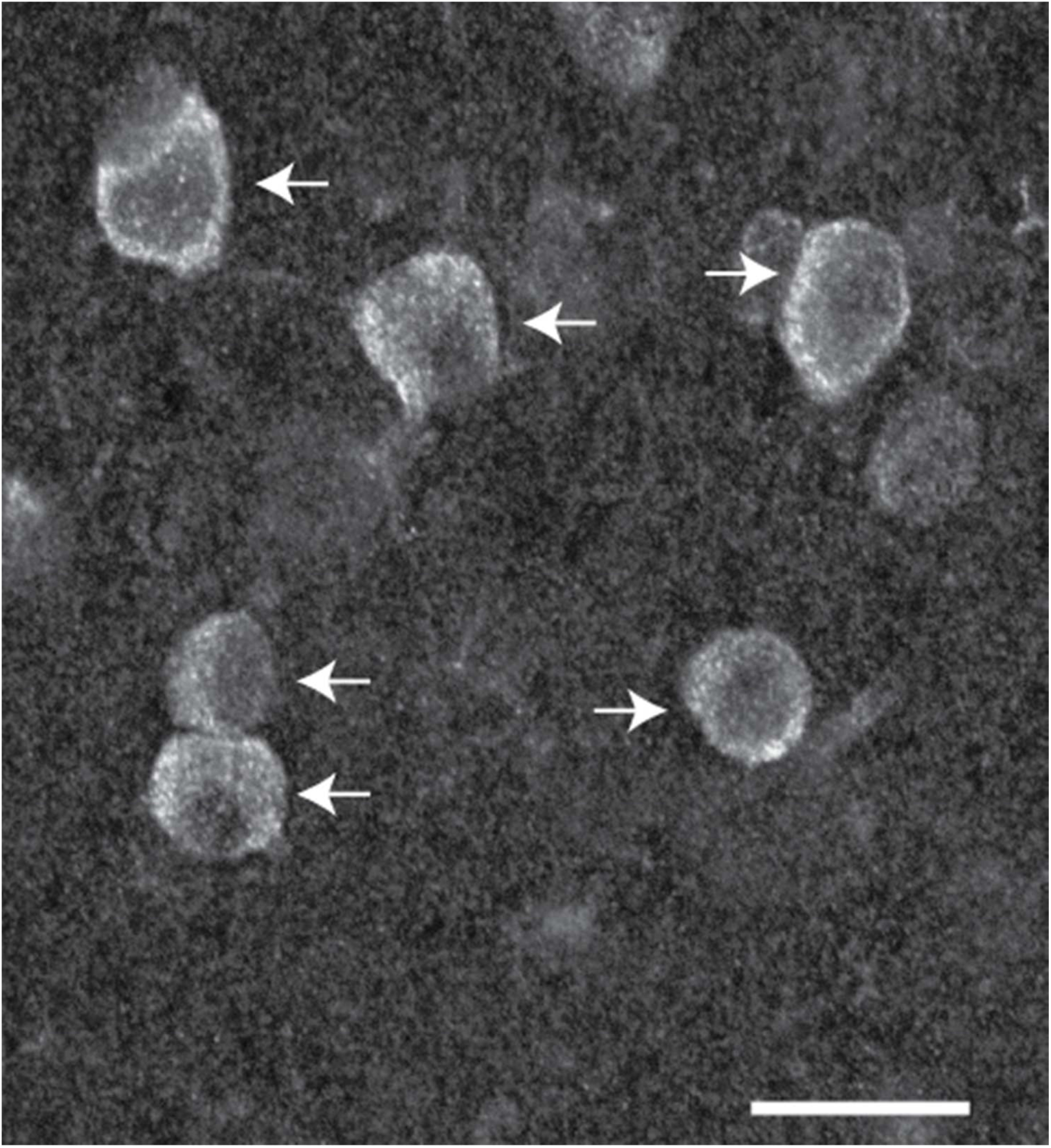
Neurons immunoreactive for the m1 acetylcholine receptor in layer III. Arrows mark cells that met counting criteria and were quantified. These neurons are characterized by strong somatic labeling appearing intense along the perimeter of the cell body. Scale bar = 20 µm.

### Dual m1AChR/CB or m1AChR/CR immunoreactivity

Because m1AChR immunoreactivity is generally restricted to the cell body and proximal dendrites, this is also the cellular compartment in which dual immunoreactivity is most commonly observed.

## Discussion

In macaque MT, we find that most CB-ir neurons express m1AChRs, while few CR-ir neurons express m1AChRs. Inhibitory interneurons immunoreactive for calcium-binding proteins exhibit structural and functional diversity. CB-ir neurons include within their population the morphological class known as the double bouquet cell, and neurogliaform and Martinotti cells have also been reported to express CB (DeFelipe, 1997). Our processing conditions yielded few CB-ir processes, making morphological classification difficult. Similar to previous reports, we found most CB-ir cell bodies to be located in layers II and III and to be interspersed among a more faintly stained CB-ir subpopulation which have been previously identified as pyramidal cells, and therefore putatively glutamatergic (Hof et al., 1999). In primates, the axons of inhibitory CB-ir neurons tend to target dendritic shafts, spines, and, to a lesser extent, somata of excitatory cells. These axons (particularly from double bouquet cells) extend vertically within a narrow cortical column, and are believed to provide inhibitory regulation of dendritic integration by excitatory neurons across layers, within a column (reviewed by DeFelipe, 1997). Further, CB-ir neurons in the dorsolateral prefrontal cortex of macaques exhibit distinct physiological properties including a long membrane time constant and action potential duration, high input resistance, and regular spiking (Zaitsev et al., 2005). It is important to note that the low-threshold spiking neuron, which in rodent cortex (at least in juveniles) expresses CB (Kawaguchi, 1995), was not observed in a study of 79 interneurons in the cortex of the adult macaque (Zaitsev et al., 2005).

The morphology of CR-ir neurons is usually described as bipolar, although double bouquet and Cajal-Retzius cell types have been identified within this population as well (DeFelipe, 1997; Meskenaite, 1997). It is not known what (if anything) distinguishes CB-ir from CR-ir double bouquet cells. We find most CR-ir neurons to be bipolar, with long, vertically oriented processes. Axonal varicosities are often observed (Figure 6 and Figure 7). These cells are also most commonly expressed in layers II and III, similar to previous reports in macaque area 17 (Meskenaite, 1997). Similar to the CB-ir population, the CR-ir population contains within it a more faintly stained subpopulation, previously identified as pyramidal (Hof et al., 1999).

Here, we report a third population of CR-ir neurons, defined by strong nuclear staining (Figure 6). Calcium-binding proteins can be expressed in the cell nucleus (Bachs and Carafoli, 1987; Gilchrist and Pierce, 1993; Nash et al., 1994). CR expression, specifically, has been reported in the nucleus of neurons in the frontal cortex of the chinchilla (Krawczyk et al., 2012). The nuclear immunoreactivity is thought to be due to the low molecular weight variant of CR (29 kDa), which (due to its smaller size) can be passively transported through the nuclear pores where it may alter gene expression (Schwaller et al., 1997). Because some of the neurons showing nuclear immunoreactivity in the present study appeared to be pyramidal in shape and because the presence of CR within the nucleus may be transient and/or activity dependent (based on its role in plastic changes in gene expression), and therefore probably does not define a stable cell class, we did not quantify this population.

The axons of CR-ir neurons, particularly those of bipolar cells, tend to target pyramidal cells in infragranular layers, but can also target other interneurons locally in the supragranular layers (DeFelipe, 1997; Meskenaite, 1997). Synapses are made onto somata, dendrites, and spines. Thus, CR-ir neurons may participate in both local (intralaminar) disinhibition, and columnar inhibition of pyramidal cells. Unlike in the cortex of rodents (Kawaguchi and Kubota, 1997), CB-ir and CR-ir neurons cannot be distinguished based on electrophysiological criteria. As such, CR-ir neurons are also observed to have long membrane time constants and action potential durations, high input resistance, and regular spiking (Zaitsev et al., 2005). Further, the burst-spiking population of CR-ir neurons that, in rodents, expresses both CR and vasoactive intestinal polypeptide (Cauli et al., 1997; Kawaguchi and Kubota, 1997; Porter et al., 1998) has not been observed in macaque (Zaitsev et al., 2005).

### m1AChR expression in V1 and MT

m1AChR expression by CR-, CB-, and PV-ir neurons in V1 and by PV-ir neurons in MT has been previously described (Disney and Aoki, 2008; Disney et al., 2014). These studies show that m1AChRs are expressed by approximately 40% of CR-ir neurons, 60% of CB-ir neurons, and 80% of PV-ir in V1 (Figure 9). In MT, expression levels are similar to V1 for PV- and CB-ir populations (75% and 55%, respectively), but the CR-ir population in MT differs from that in V1 (10% compared to 40%, respectively). Because the levels of m1AChR expression we observe in PV-ir and CB-ir neurons are comparable in V1 and MT, and because the overall immunoreactivity appears similar in both areas, it is unlikely that the observed difference in expression by the CR-ir population is due to a detection failure.

**Figure 9:**
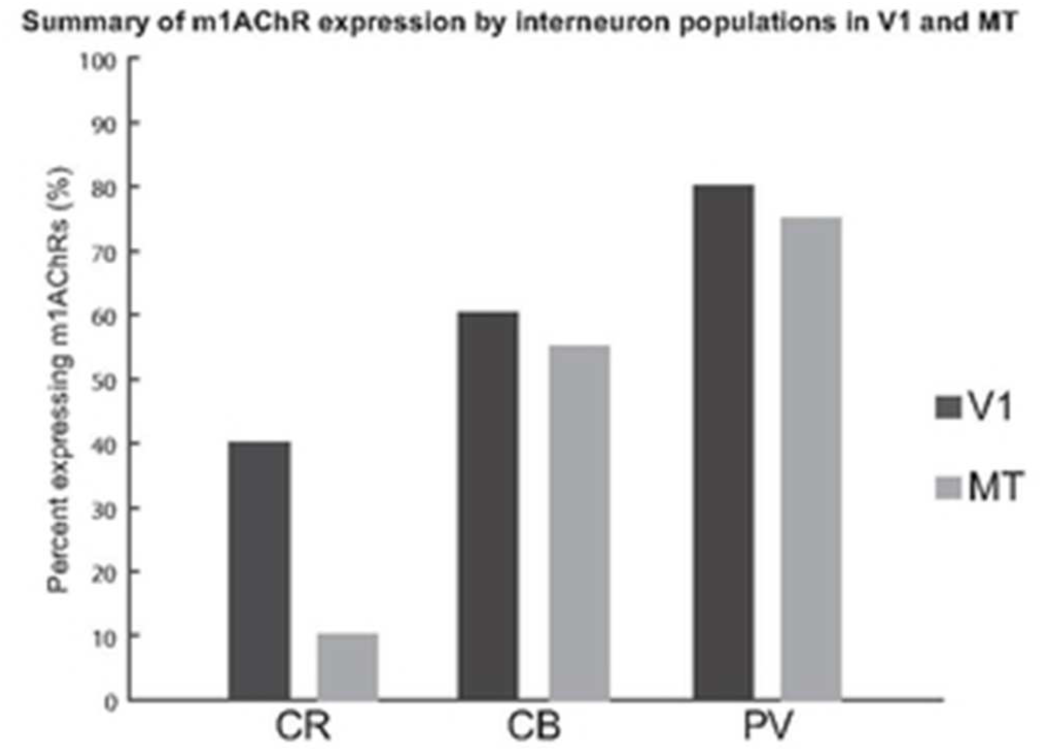
Comparison of the m1 acetylcholine receptor (m1AChR) expression by populations immunoreactive for calretinin (CR), calbindin-D28k (CB), and parvalbumin (PV) in primary visual area V1 (dark gray) and middle temporal area MT (light gray). In V1, 40% of CR-immunoreactive neurons, 60% of CB-immunoreactive neurons, and 80% of PV-immunoreactive neurons express m1AChR. Those numbers in MT are 10%, 55%, and 75%, respectively.

If the lower level of m1AChR expression by CR-ir neurons in MT is a characteristic of areas in the dorsal visual pathway (of which MT is a part), we would predict that the V1 layer that projects to MT (layer 4B) might also show similarly low expression levels by the CR-ir neuronal population. CR is not often expressed by neurons in layer 4B; Disney et al. (2008) found only seven CR-ir neurons in layer 4B across three animals. Of those seven, four expressed m1AChRs. While the number of reported cells is sparse, this does not appear support a hypothesis that low m1AChR expression by the CR-ir population is a feature of the dorsal stream. In this context, it will be interesting to investigate m1AChR expression by PV-, CB-, and CR-ir neurons in visual area V4, which represents an area within the ventral visual stream (the ‘what’ pathway) at roughly the same hierarchical level as MT and in later stages of the “where” pathway such as the lateral intraparietal area. Perhaps low m1AChR expression by the CR-ir population represents a feature mid-level visual processing. These studies are currently underway in our lab.

It is also possible that CR-ir neurons in area MT express one of the other Gq-coupled muscarinic ACh receptors. The m3 and m5 muscarinic receptors are members of the same pharmacological class as the m1AChR (M1) and have the same G protein-coupling, but different affinities for ACh than does the m1AChR. Unfortunately, at this time, the possibility that another Gq-coupled receptor type has been substituted in CR-ir neurons in MT cannot be addressed using similar methods, as there are currently no antibodies directed against the m3 or m5 muscarinic ACh receptors that pass controls for use in macaques. Studies to instead address this question by *in situ* hybridization of mRNA transcripts for the m1, m3, and m5 receptors are underway in our lab.

### Calcium-binding protein immunoreactive neurons as cholinergic targets

As demonstrated here and in previous studies (Disney and Aoki, 2008; Disney et al., 2014), neurons immunoreactive for the calcium-binding proteins PV, CB, and CR represent targets for cholinergic modulation in the visual cortex. It has been hypothesized that ACh release supports cortical processing during various attentive states (Sarter et al., 2005; Herrero et al., 2008), potentially implicating inhibitory interneurons in attentive processes. The biased competition model for attention (Desimone et al., 1990; Desimone and Duncan, 1995; Desimone, 1998) proposes that neuronal representations of stimuli compete for further processing and that this competition can be biased by top-down and/or bottom-up processes. This bias represents one possible mechanism by which preferential processing can be conferred at a circuit level upon “attended” objects or locations. The ability to control columnar inhibition has clear relevance for a biased competition model in an area, like MT, in which properties such as position in visual space and direction of motion are mapped in a systematic fashion across the cortical sheet.

If ACh release does support attentive states, then the fact that ACh receptor expression differs between V1 and MT implies that this biased competition would be expected to play out differently in the two areas. CR-ir neurons in V1 can both provide a local disinhibition within the superficial layers and columnar inhibition of pyramidal cells in deeper layers (Meskenaite, 1997). However, we are hesitant to use what is known about CR-ir targeting in V1 as a proxy for CR-ir regulation in MT, as inhibitory interneuron populations differ even between visual areas V1 and V2 in macaques (DeFelipe et al., 1999). As such, specific data on the laminar profile of post-synaptic targets for inhibition mediated by CR-ir neurons in MT are needed. Once these data are available, it is likely that computational modeling of modulated cortical circuits will be needed to yield predictions regarding the form of the resulting modulatory differences between these two functionally distinct cortical areas/compartments.

### Neuromodulatory compartments

Differences in local anatomical features (such as differences in receptor expression by cell type, shown here) provide a mechanism for a variation in cholinergic modulation across cortical areas. We have previously proposed the existence of unique neuromodulatory compartments across cortex (Coppola et al., 2016), that are determined by anatomical characteristics. Here, we report differences in modulatory compartments in cortex that align with functional compartmentalization in vision (i.e. V1 versus MT). The clear implication of these data is that, even if the cholinergic signal delivered to V1 and MT is the same, the circuit response to that modulation will differ. This is one of many possible scales on which the modulatory compartmentalization of cortex exists—some are finer grained (such as laminar differences in acetylcholinesterase expression) and others will be coarser (such as differences between frontal and occipital cortices in overall innervation density). Regardless of the scale, a nuanced understanding of the structural determinants of cholinergic function and how they differ both between and within species (Coppola and Disney, Under review) will be essential if we are to understand when and how neuromodulators contribute to the dynamical control of cortical state.

## Acknowledgments

The authors thank the Vanderbilt University Cell Imaging Shared Resource for technical support and facilitation of confocal imaging (supported by NIH grants CA68485, DK20593, DK58404, DK59637, EY08126, and 1S10RR027396-01). We also thank Juliane Krueger for assistance with data analysis. This work was supported by NIH grant R00 MH-93567 (AAD).

